# Frailty is related to serum inflammageing markers: results from the VITAL study

**DOI:** 10.1101/2023.08.24.554610

**Authors:** Yannick van Sleen, Sudarshan A Shetty, Marieke van der Heiden, Milou CA Venema, Nicolas Gutierrez-Melo, Erik JM Toonen, Josine van Beek, AnneMarie M Buisman, Debbie van Baarle, Delphine Sauce

**Affiliations:** University Medical Center Groningen, Groningen, The Netherlands; Sorbonne Université, INSERM, Centre d’Immunologie et des Maladies Infectieuses, Cimi-Paris, F-75013, Paris, France; R&D Department, Hycult Biotechnology, Uden, the Netherlands; Center of Infectious Diseases Control, National Institute of Public Health and the Environment, Bilthoven

**Keywords:** Ageing, Frailty, Inflammation, Biomarkers, Body Mass Index, Cytomegalovirus

## Abstract

Frailty describes an age-associated state in individuals with an increased vulnerability and less resilience against adverse outcomes. To score frailty, studies have employed the questionnaires, such as the SF-36 and EQ-5D-3L, or the Frailty Index, a composite score based on deficit accumulation. Furthermore, ageing of the immune system is often accompanied by a state of low-grade inflammation (inflammageing). Here, we aimed to associate 29 circulating markers of inflammageing with frailty measures in a prospective cohort study to understand the mechanisms underlying ageing.

Frailty measures and inflammageing markers were assessed in 317 participants aged 25-90. We determined four different measures of frailty: the Frailty Index based on 31 deficits, the EQ-5D-3L and two physical domains of the SF-36. Serum/plasma levels of inflammageing markers and CMV/EBV seropositivity were measured using different techniques: Quanterix, Luminex or ELISA.

All four measures of frailty strongly correlated with age and BMI. Nineteen biomarkers correlated with age, some in a linear fashion (IL-6, YKL-40), some only in the oldest age brackets (CRP), and some increased at younger ages and then plateaued (CCL2, sIL-6R). After correcting for age, biomarkers, such as IL-6, CRP, IL-1RA, YKL-40 and elastase, were associated with frailty. When corrected for BMI, the number of associations reduced further.

In conclusion, inflammageing markers, particularly markers reflecting innate immune activation, are related to frailty. These findings indicate that health decline and the accumulation of deficits with age is accompanied with a low-grade inflammation which can be detected by specific inflammatory markers.

## INTRODUCTION

There are major differences in how individuals and their immune systems cope with ageing. Ageing of the immune system leads to a number of profound changes to the composition of leukocyte subsets and their functioning (1). Older age has been associated with increased chronic inflammation, as evidenced by elevated C-reactive protein (CRP), TNF-α and IL-6 levels; a process also known as inflammageing (2–4). Inflammageing is typically characterized by an enhanced innate immune response (5), evidenced by increased circulation of proteins associated with macrophages (e.g. soluble CD163 (sCD163), YKL-40, sCD14) and neutrophils (e.g. elastase, PR3, IL-8). Adaptive cells are also vulnerable to the ageing process; their diversity decreases substantially with age. Partly, this has been associated with increased exposure to chronic viral infections such as cytomegalovirus (CMV) and Epstein-Bar virus (EBV), resulting in the expansion of these antigen-specific memory cells also known as memory inflation (6,7).

The term ‘frailty’ is used to describe older adults with a declining health and signifies an increased vulnerability to adverse outcomes, most notably: physical impairment, disease and mortality. Frailty largely concerns the ability of an organism to cope with outside stressors, and is found more often in those with comorbidities and in those in long-term care (8–11). It is a dynamic state which tends to increase with age, but can also fluctuate over time. Moreover, frailty has also been associated to the body mass index (BMI), with higher frailty scores in people with a too low or too high BMI (12). Understanding the mechanisms behind frailty is of high relevance since the population is aging worldwide.

There are a number of approaches to measure frailty. A frequently used method to determine an individual’s frailty, is by the Frailty Index as described by Rockwood and Mitnitski, which is known to be a better predictor of hospitalization and mortality than age by itself (13,14). The Frailty Index signifies the accumulation of an individual’s deficits ranging from impaired hearing to the presence of heart failure. Another important instrument to determine this phenomenon, the Frailty Phenotype, has been developed by Fried et a*l*., (2001) which measures frailty based on a number of tests such as the grip strength. Other, more subjective methods measure the quality of life using on questionnaires, such as the EuroQol-5 Dimensions-3 Level (EQ-5D-3L) and Short-form (SF)-36 (15,16). These methods are often employed to assess patient-reported outcomes after interventions, for example during clinical trials, although aspects of these questionnaires are also used as measures of frailty (17).

Considering the accumulation of deficits in frail individuals, we hypothesize that inflammageing occurs more extensively in frail than non-frail individuals, and that there are specific markers that reflect these differences the best. Therefore, in this study, we aimed at associating frailty measures and serum markers of inflammageing in participants of three different age groups (adults, middle-aged and older adults) of the VITAL study (*Vaccines and Infectious disease in the Ageing Populations*) (18,19). To this end, we selected a number of markers that were previously associated with frailty (e.g. IL-6, CRP, neopterin, iFABP2) and supplemented these with a number of markers associated with systemic inflammation and innate immunity (2–4,20–24). We investigated the associations between these markers and four measures of frailty: the Frailty Index, the EQ-5D-3L, and two domains of the SF-36.

## RESULTS

### Frailty is strongly associated with age and BMI

Frailty measures were documented in all participants divided over three age groups, with the oldest age group (≥65 years of age) containing the most participants (Table 1). Participants in the younger age group (25-49 years of age) were more often female than participants in the older groups. The median age of male participants was 7 years higher than that of female participants (p=0.014; Figure 1A). We therefore corrected for sex when assessing the effect of age on frailty measures.

**Table 1:** characteristics of VITAL cohort participants included in this study.

|  | **Total** | **Young**<br>**(25-49 years)** | **Middle age**<br>**(50-64 years)** | **Older**<br>**(≥65 years)** |
| --- | --- | --- | --- | --- |
| **N** | 3 | 63 | 95 | 159 |
| **Age** median (IQR) | 65 (54-75) | 35 (29-42) | 58 (55-61) | 75 (69-81) |
| **Sex** % female | 54 | 67 # | 59 | 47 |
| **BMI** kg/m<sup>2</sup>, median (IQR) | 24.6<br>(22.4-27.5) | 24.0 #<br>(21.5-25.5) | 24.0 &<br>(21.8-26.5) | 25.3<br>(23.4-28.6) |
| **CMV** %positive / %negative | 48/51 | 42/57 | 51/47 | 50/48 |
| **EBV** %positive / %negative | 83/14 | 76/18 | 87/13 | 83/13 |
| **Frailty index** median (IQR) | 0.13<br>(0.08-0.20) | 0.06 # \$<br>(0.03-0.10) | 0.10 &<br>(0.06-0.14) | 0.18<br>(0.13-0.26) |
| **EQ-5D-3L** median (IQR) | 1 (0.81-1) | 1 # (0.84-1) | 1 & (0.84-1) | 0.84 (0.81-1) |
| **SF-36 PF** median (IQR) | 95 (80-100) | 100 # \$ (95-100) | 95 & (90-100) | 85 (65-95) |
| **SF-36 HG** median (IQR) | 70 (60-80) | 75 # (70-85) | 75 & (65-85) | 65 (55-75) |
| **Health rating** median (IQR) | 8.0<br>(7.0-8.5) | 8.0<br>(7.5-9.0) | 8.0<br>(7.5-9.0) | 8.0<br>(7.0-8.5) |
| **Number of medications** median (IQR) | 2 (0-4) | 0 # (0-1) | 1 & (0-2) | 4 (2-6) |

**Figure 1:**
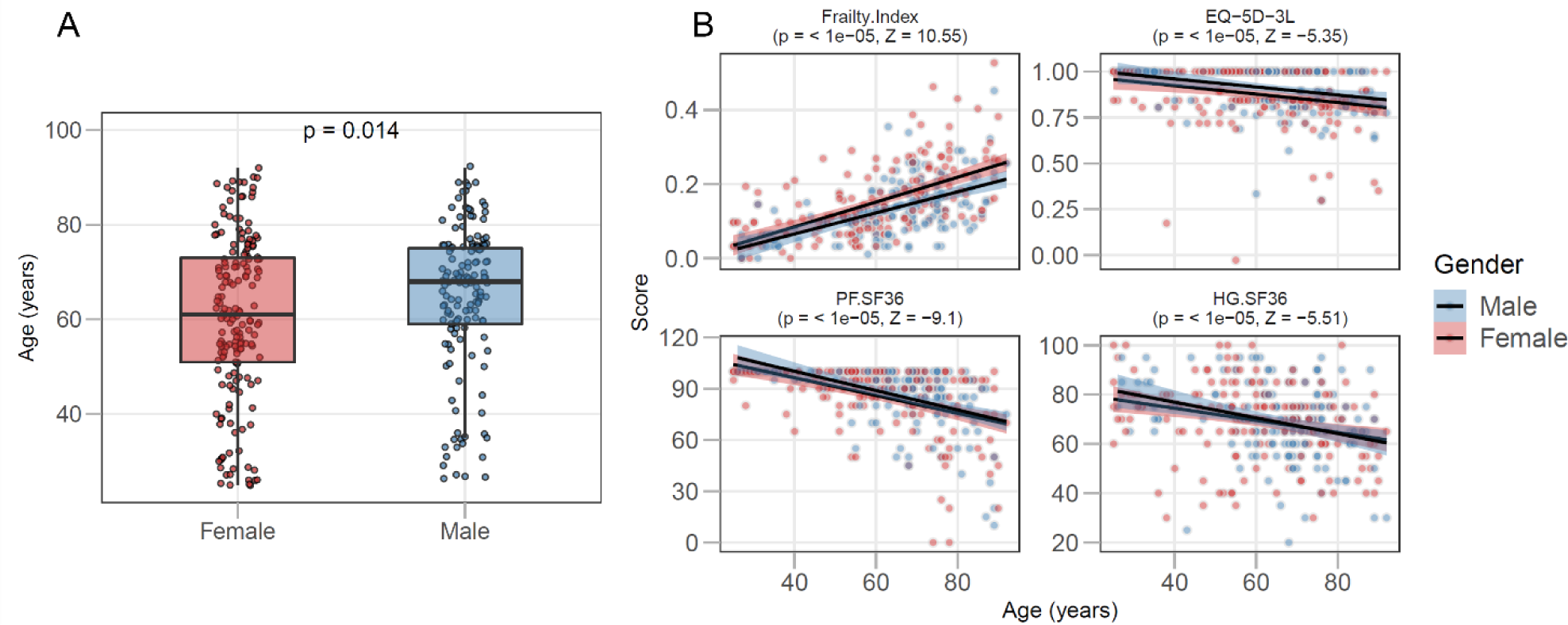
Measures of frailty are significantly associated with age. Graph A shows the age distribution of the total study population (n=317) by sex. In B, shown are associations of participant age with the Frailty Index, the EQ-5D-3L (EuroQol-5 Dimensions-3 Level) questionnaire, the PF (physical functioning) and HG (general health) domains of the SF-36 (short-form-36). Strength and significance of the sex-corrected Spearman correlation coefficient is indicated in each graph. The regression line is shown in black and the shaded region represents the confidence interval.

The Frailty Index, scores of the EQ-5D-3L questionnaire and scores in the physical functioning (PF) and general health (HG) domains of the SF-36 correlated significantly with age, when corrected for sex (p<0.0001; Figure 1B). For the Frailty Index, we observed an accumulation of deficits with increasing age, with the natural logarithm of the slope of the mean at 0.029, which corresponds with previous reports that used this index (27,29). However, the highest Frailty Index score found within our study was 0.53, which is slightly lower than the defined submaximal limit of 0.67 (13,14). Measures of frailty correlated with each other, with the strongest association between the Frailty Index and the number of prescription medications (Spearman’s R=0.70, Supplementary Figure S1A).

Frailty measures were also associated with BMI (Supplementary Figure S1B). Participants with a higher BMI tended to be frailer than participants with a BMI lower than 25. In our study population, male participants had a significantly higher BMI than females (p=0.006). However, BMI also correlated positively with age (p= 0.003, Z-statistic=2.96) after controlling for sex.

Participants that were seropositive for CMV, but not EBV, had a significantly higher Frailty Index score (p=0.048). Age only had a moderate effect on CMV and EBV status, particularly in participants aged over 50 years (Table 1).

### Changes in concentration of serum inflammageing markers follow different trajectories with age

Out of 29 biomarkers measured, 19 of them were significantly associated with age, when corrected for sex (Table 2, Supplementary Figure S2). The patterns of levels of inflammageing markers were however not uniform: four different trajectories could be identified (Figure 2A). The levels of six biomarkers increased steadily from young to middle aged to older adults (trajectory a). In the second trajectory, we observed elevated levels of five markers (e.g. CCL2) in middle aged and older adults compared to younger adults, but no further increase in adults over 65 years (trajectory b). The third trajectory is characterized by elevated levels in the older adults only, which is observed for eight markers (e.g. PR3; trajectory c). The final category of ten serum markers was not associated with age (trajectory d).

**Table 2:** levels of inflammageing markers differ between age groups.

| Median (IQR) | Unit | Total | Young | Middle-age | Older |
| --- | --- | --- | --- | --- | --- |
| **N** |  | 317 | 63 | 95 | 159 |
| **Group a** |  |  |  |  |  |
| **IL-6** | pg/mL | 0.89<br>(0.46-1.37) | 0.40<br>(0.22-0.84) | 0.69<br>(0.43-1.12) | 1.10<br>(0.72-1.74) |
| **Elastase** | ng/mL | 208<br>(153-296) | 175<br>(128-220) | 195<br>(154-271) | 228<br>(166-338) |
| **sCD163** | ng/mL | 745<br>(518-1134) | 529<br>(381-547) | 681<br>(530-1039) | 906<br>(627-1244) |
| **CXCL10 (IP-10)** | pg/mL | 24.7<br>(18.8-31.7) | 18.8<br>(14.7-23.2) | 21.2<br>(17.5-28.5) | 29.1<br>(22.1-35.4) |
| **YKL-40** | ng/mL | 37.6<br>(26.0-67.7) | 24.3<br>(19.7-31.2) | 30.3<br>(22.0-42.9) | 58.2<br>(35.8-93.6) |
| **Neopterin** | nmol/L | 8.47<br>(5.74-11.33) | 5.78<br>(4.11-8.01) | 7.65<br>(5.64-9.72) | 10.45<br>(7.13-13.95) |
| **Group b** |  |  |  |  |  |
| **CCL2** | pg/mL | 402<br>(317-498) | 335<br>(290-416) | 413<br>(319-519) | 417<br>(339-523) |
| **IL-8** | pg/mL | 20.5<br>(14.8-34.1) | 14.9<br>(11.5-21.4) | 20.5<br>(15.3-42.6) | 22.3<br>(16.4-35.4) |
| **sIL-6R** | ng/mL | 46.9<br>(41.3-52.9) | 42.8<br>(37.3-49.6) | 47.1<br>(41.3-52.9) | 48.4<br>(42.4-53.8) |
| **iFABP2** | pg/mL | 1138<br>(843-1644) | 908<br>(660-1233) | 1113<br>(863-1507) | 1235<br>(932-1839) |
| **sGP130** | ng/mL | 143<br>(131-157) | 134<br>(120-145) | 143<br>(130-158) | 147<br>(136-158) |
| **Group c** |  |  |  |  |  |
| **PR3** | ng/mL | 32.4<br>(24.5-46.8) | 27.9<br>(22.6-38.6) | 28.7<br>(19.9-42.9) | 37.9<br>(26.4-52.7) |
| **CRP** | mg/L | 1.19<br>(0.55-2.82) | 0.75<br>(0.23-1.89) | 1.00<br>(0.45-2.29) | 1.62<br>(0.68-3.31) |
| **sCD14** | ng/mL | 1446<br>(1252-1645) | 1327<br>(1129-1533) | 1418<br>(1229-1573) | 1661<br>(1135-2254) |
| **Angiopoietin-2** | pg/mL | 2023<br>(1593-2742) | 1787<br>(1268-2545) | 1909<br>(1545-2319) | 2233<br>(1749-3000) |
| **sCD25** | pg/mL | 522<br>(427-653) | 467<br>(381-547) | 478<br>(388-61) | 595<br>(488-718) |
| **IL-1RA** | pg/mL | 855<br>(670-1106) | 708<br>(567-975) | 770<br>(651-992) | 936<br>(779-1245) |
| **IFN-γ** | pg/mL | 0.04<br>(0.03-0.07) | 0.03<br>(0.02-0.04) | 0.04<br>(0.02-0.06) | 0.05<br>(0.03-0.08) |
| **SAA** | ng/mL | 135<br>(6-735) | 45<br>(5-356) | 67<br>(5-356) | 259<br>(39-886) |
| **Group d** |  |  |  |  |  |
| **Calprotectin** | ng/mL | 1586<br>(1102-2310) | 1538<br>(772-2084) | 1661<br>(1135-2254) | 1661<br>(1135-2254) |
| **C5a** | ng/mL | 21.3<br>(16-28.6) | 19.5<br>(15.6-25.5) | 19.6<br>(14.7-28.7) | 22.7<br>(16.9-29.5) |
| **PTX-3** | pg/mL | 5519<br>(4103-7188) | 5328<br>(3789-6988) | 5402<br>(4032-6949) | 5679<br>(4697-7278) |
| **Cathepsin G** | ng/mL | 6.62<br>(4.70-8.66) | 6.48<br>(4.60-8.04) | 6.10<br>(4.65-8.83) | 7.20<br>(4.83-8.77) |
| **A1AT/elastase complex** | µg/mL | 56<br>(38-96) | 55<br>(9-106) | 73<br>(26-135) | 52<br>(39-81) |
| **GM-CSF** | pg/mL | 0.17<br>(0.13-0.22) | 0.17<br>(0.13-0.23) | 0.17<br>(0.12-0.22) | 0.18<br>(0.13-0.22) |
| **IL-1β** | pg/mL | 0.03<br>(0.01-0.6) | 0.02<br>(0.01-0.05) | 0.04<br>(0.02-0.07) | 0.03<br>(0.01-0.06) |
| **TNF-α** | pg/mL | 2.22<br>(1.87-2.68) | 2.08<br>(1.72-2.52) | 2.31<br>(1.81-2.68) | 2.21<br>(1.81-2.68) |
| **IL-10** | pg/mL | 0.57<br>(0.42-0.83) | 0.50<br>(0.38-0.68) | 0.66<br>(0.45-0.86) | 0.56<br>(0.40-0.84) |
| **IFN-α** | pg/mL | 0.01<br>(0.01-0.02) | 0.01<br>(0.01-0.02) | 0.01<br>(0.00-0.02) | 0.01<br>(0.01-0.02) |

Markers are sorted per group depending on the pattern as seen in Figure 1. Shown is median +/- IQR. IQR: inter-quartile range.

**Figure 2:**
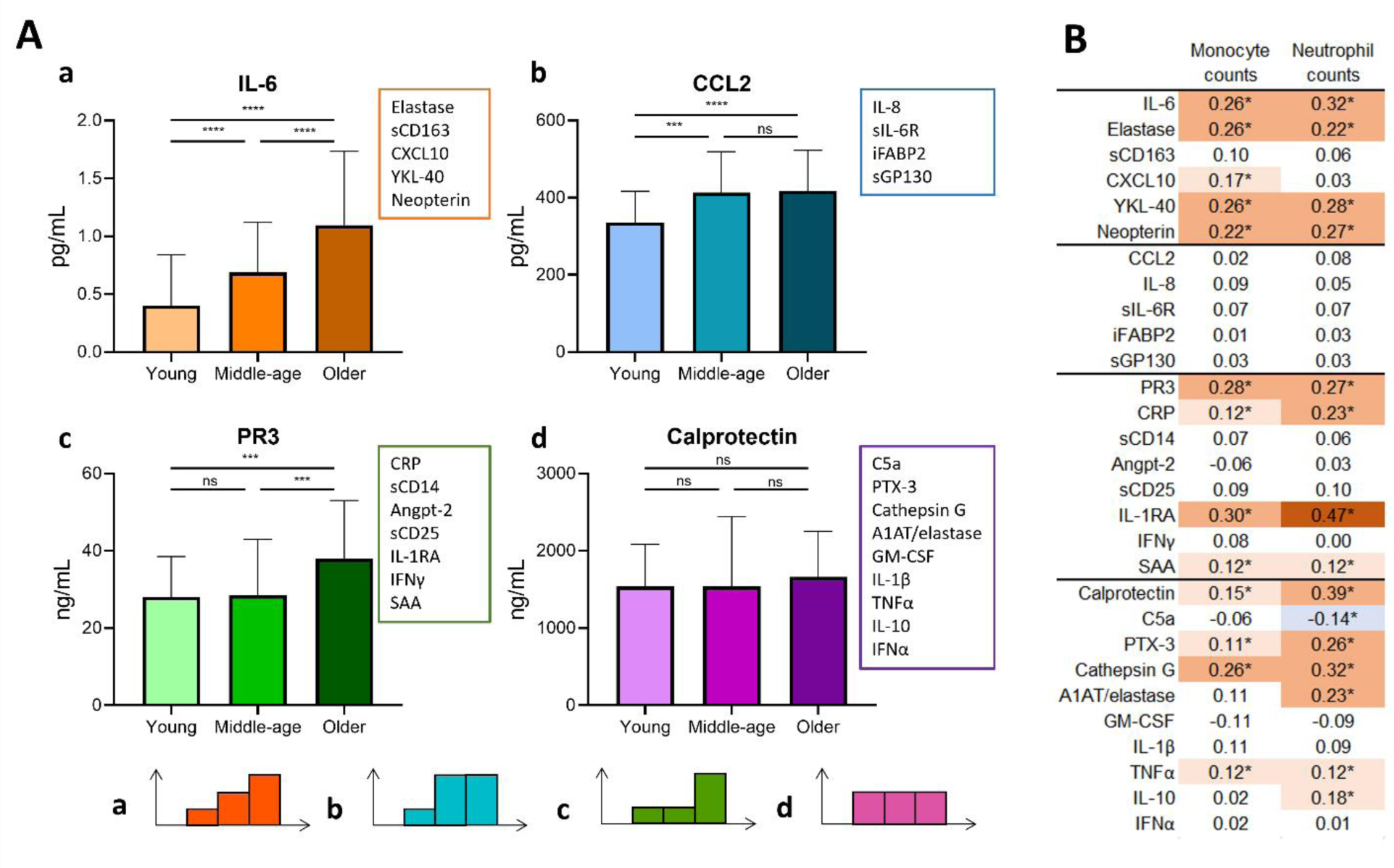
Trajectories of serum markers of inflammageing in three age groups and their association with innate cells. In A, four different trajectories could be identified, based on statistical significance (Mann Whitney U) between the groups. Shown are an example of each trajectory and the other markers that follow this pattern. #: these markers were not identified in the sex-corrected analysis as markers that significantly associate with age, but did follow the same trajectory, based on statistical significance, as the other markers in their list. In B, we show the correlations of circulating monocyte and neutrophil counts with the biomarkers. Shown are the Spearman R coefficients, with color coding according to the strength of the association. *: p<0.05, *** : p<0.001, **** : p< 0.0001, ns: not significant.

As many of the biomarkers that we selected were associated with activation of cells of the innate immune system, we examined whether absolute counts of circulating monocytes or neutrophils correlated with the serum/plasma levels of the biomarkers (Figure 2B). Interestingly, we show that a number of markers in trajectory a, c and d, but not in trajectory b, associate with monocyte and/or neutrophil counts. The strongest associations were seen for IL-1RA, whose levels correlated with monocyte counts (Spearman R=0.30) and neutrophil counts (R=0.47).

As chronic infections with CMV and EBV have been associated with ageing of the immune system (6,30), we investigated their association with the markers of inflammageing stratified by age and sex (Supplementary Figure S3). Serum levels of the pro-angiogenic protein angiopoietin-2 showed a positive association with both CMV (p= 0.01861, Z = 2.35) and EBV (p= 0.0070, Z= 2.66) positivity, independent of age and sex. In addition, sCD163 levels were found to be higher in CMV-positive participants (p= 0.00048, Z= 3.49), and CCL-2 levels in EBV-positive participants (p= 0.01611, Z= 2.39). For all other markers, no significant association was observed.

### Markers of inflammageing associate with frailty in older participants

To explore the interplay between inflammageing and frailty, we correlated the frailty measures in the VITAL cohort with the biomarker levels. First, we associated all biomarkers with the Frailty Index using a principle component analysis (PCA) in the participants over 60 years of age, because above this age frailty measures have higher relevance than in younger people (Figure 3). We then colored the individuals based on their Frailty Index score, and show that individuals with higher scores cluster differently than individuals with lower Frailty Index scores. We next aimed to determine which biomarkers are the main drivers of this differential clustering.

**Figure 3:**
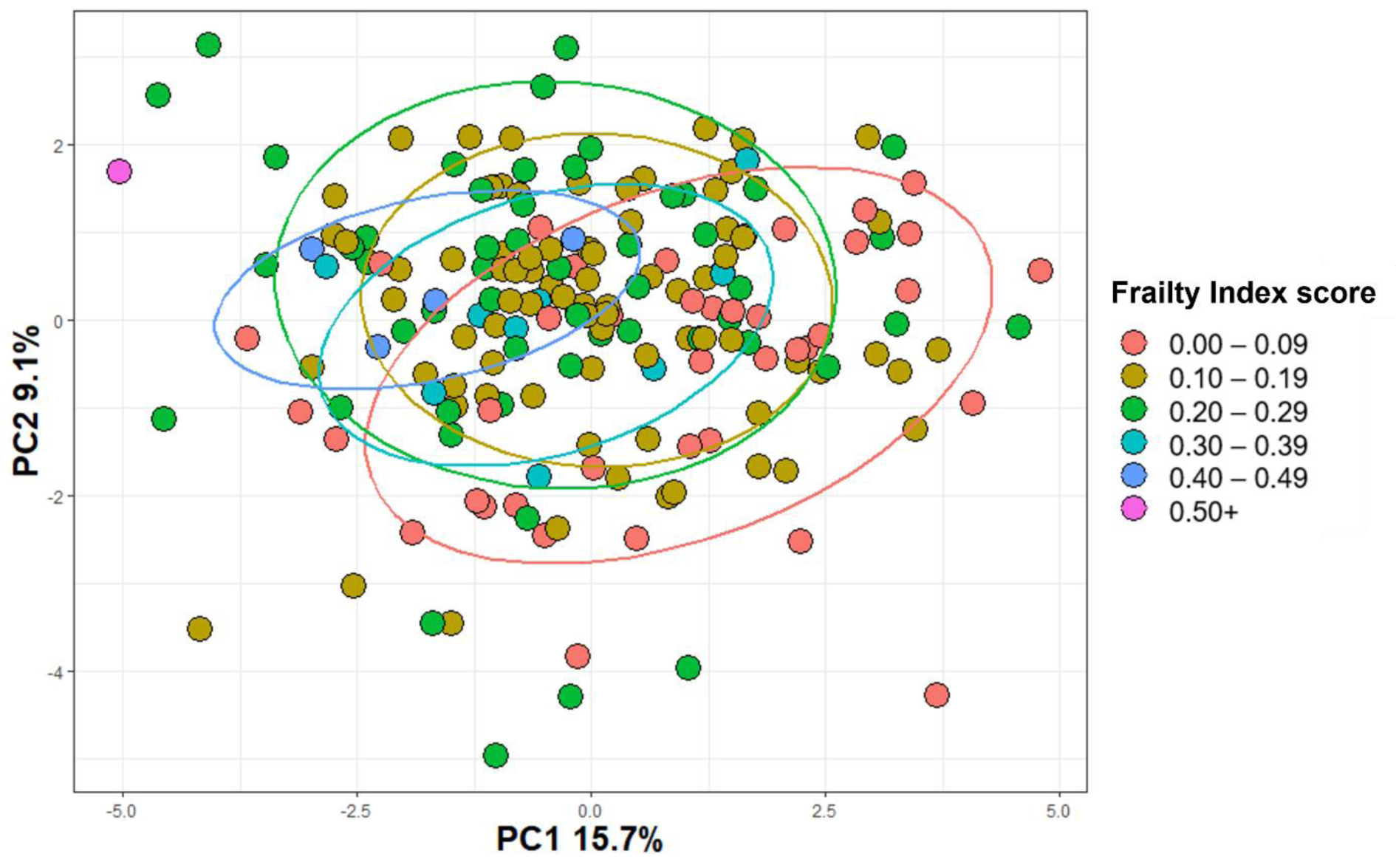
Principal component analysis (PCA) comparing the inflammageing markers in participants over 60 years of age between different classes of frailty scores. The log-transformed markers were included in the PCA analysis. Participants (n=187) were divided into different frailty classes; the red dots represent the participants with the lowest frailty scores and the blue and purple dots the participants with the highest frailty scores. The percentage of variance explained by the Principle Components (PC) 1 and 2 are mentioned on the axis. The eclipses cover 80% of the datapoints.

To this end, we first analysed which of the inflammageing markers associated with frailty as an absolute measure in all participants (Figure 4A). Out of 29 markers, 21 showed an association with at least one frailty measure (the Frailty Index, EQ-5D-3L and SF-36 domains PF and HG). Even though this implies a relation between markers of inflammageing and frailty, we already showed that age has a strong association with levels of these markers.

**Figure 4:**
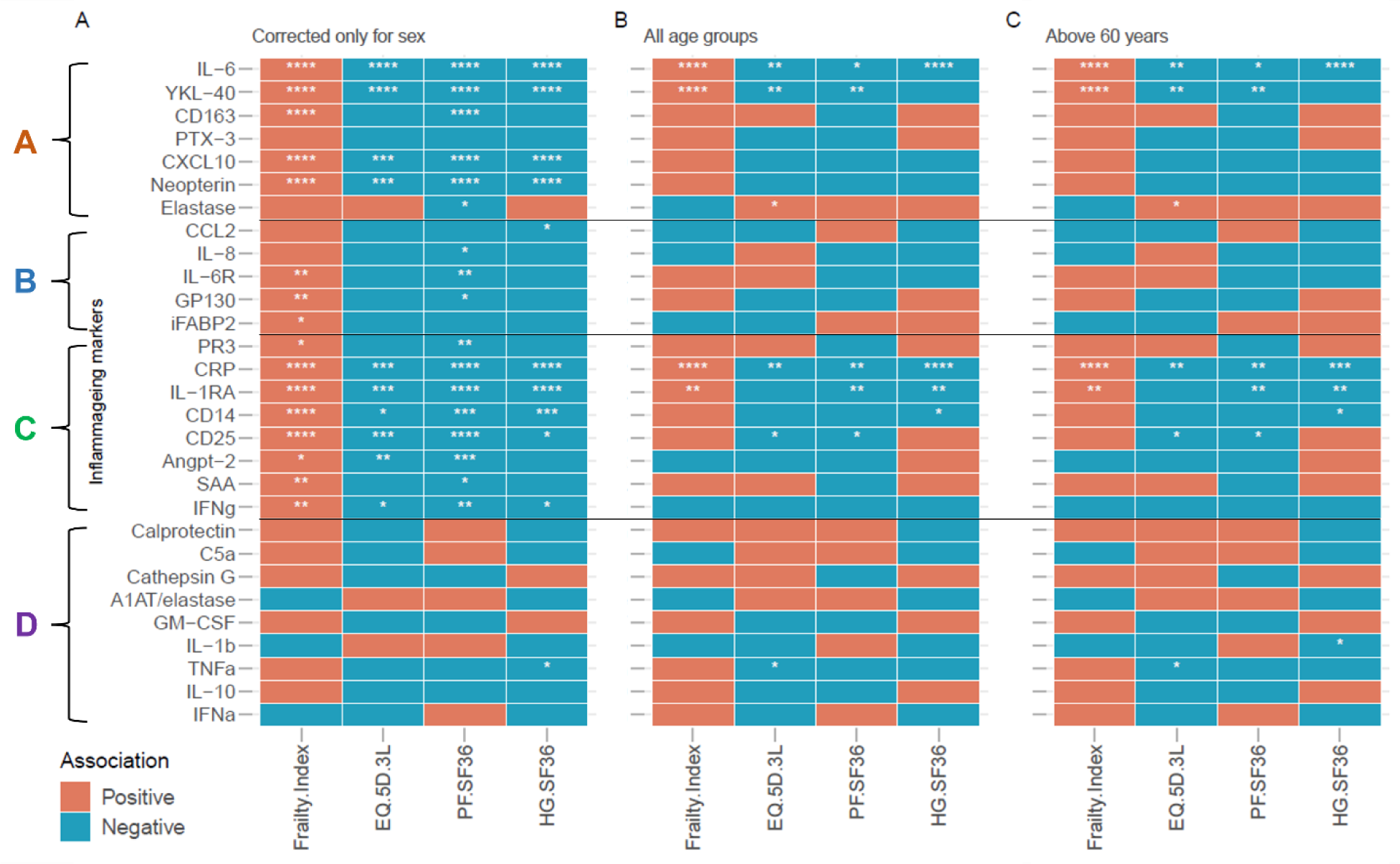
Frailty measures are associated with elevated levels of inflammageing markers. Here, we show the correlation of 29 markers with the outcomes of frailty in participants in the VITAL study (n=317). In A we show that inflammageing markers were associated with frailty as an absolute measure, only corrected for sex. In B, we show the age and sex-corrected correlations of the markers with the outcomes of frailty. In C, we show age- and sex-corrected associations in 60 plus participants. The inflammageing markers are divided in groups A, B, C, and D based on their trajectory with age, see Figure 2. Statistical significance of the Spearman’s correlation coefficient is indicated with *: p<0.05, **: p<0.01, ***: p<0.001, ***: p<0.0001. EQ-5D-3L: (EuroQol-5 Dimensions-3 Level), PF (physical functioning), HG (general health), SF-36 (short-form-36). The effect of age and sex was controlled in univariate correlation analyses using the blocked Spearman rank tests as implemented in the R package coin (v1.4.2) was used with the distribution parameter approximate (nresample=100000).

Next, we aimed to identify markers that associate with frailty in individuals of the same age. Therefore, we analyzed the association of the different frailty measures with markers of inflammageing in all VITAL cohort participants correcting for age and sex as confounders (Figure 4B). The Frailty Index was associated with four, EQ-5D-3L with six, PF.SF36 with five and HG.SF36 with four inflammageing markers. IL-6 and CRP were associated with all four frailty measures, and innate immune markers YKL-40 and IL-1RA associated with three out of the four frailty measures. The direction of these associations was the same for all markers: a positive correlation with the Frailty Index and a negative correlation with the scores on the EQ-5D-3L and SF-36 domains. The only exception was Elastase, which correlated positively with EQ-5D-3L scores. Additionally, we analysed the associations in the individuals over 60 years, and showed that this analysis did not lead to substantially different outcomes (Figure 4C).

We additionally confirmed the predictive value of the biomarkers and age on the Frailty Index using a regression analysis (Table 3). As expected, age was the strongest predictor of the Frailty Index. Additionally, CRP and IL-1RA were discovered as independent predictors of the Frailty Index. We next analyzed the associations separately for male and female participants. Besides age, IL-6 and IL-1RA levels independently predicted the Frailty Index in female participants. In male participants, YKL-40 and PR3 levels were found as independent predictors, with YKL-40 levels positively and PR3 levels negatively associated with the Frailty Index. Thus, this analysis confirms the predictive values of the inflammageing biomarkers (especially CRP, IL-6, IL-1RA and YKL-40) for frailty and reveals sex differences in this prediction.

**Table 3:**
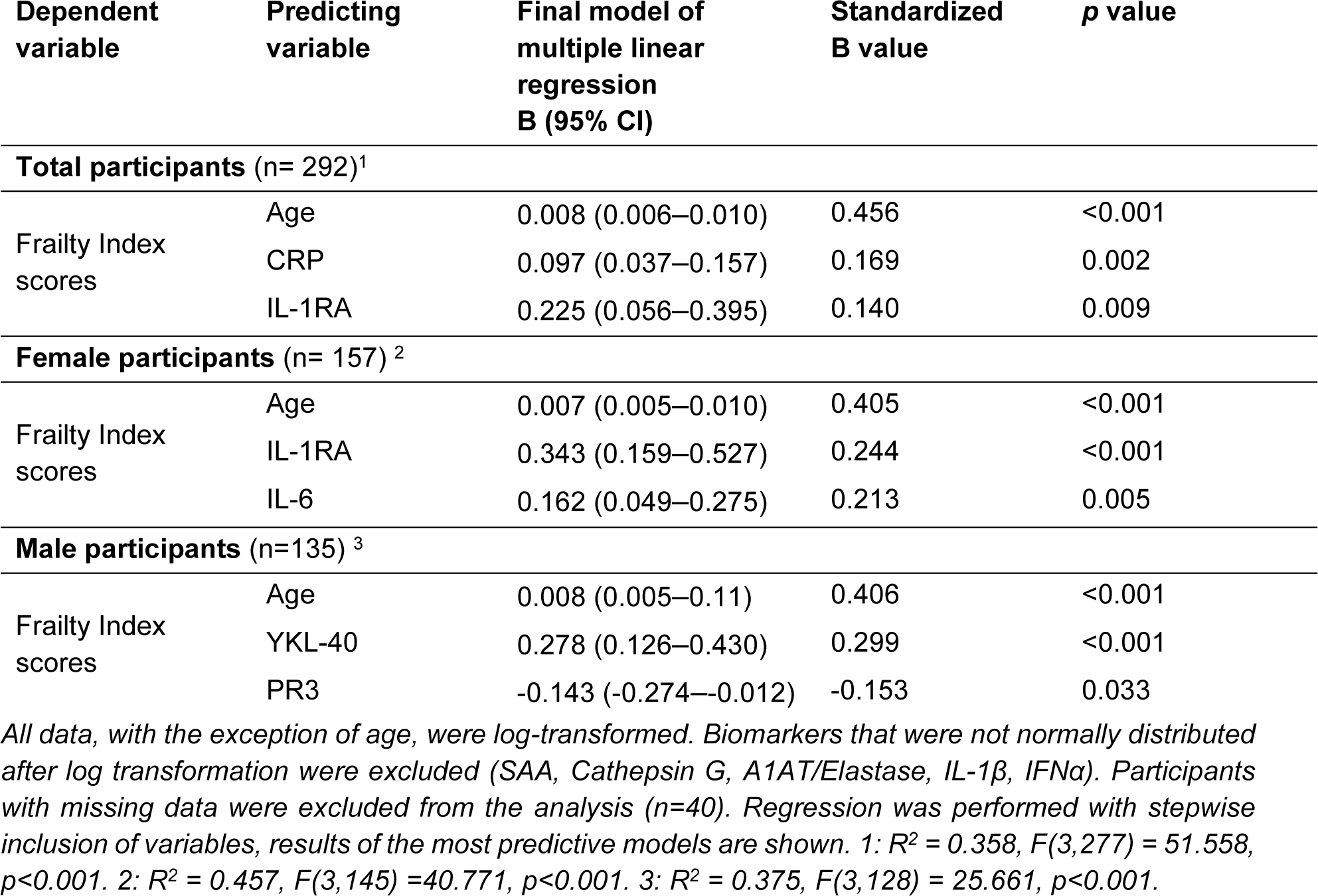
Multiple linear regression analysis of associations of the inflammageing markers and age.

### BMI may partly influence the association between frailty and inflammageing markers

Both CRP and IL-6 have previously been associated with BMI and visceral adiposity (32–34), and in our cohort we observe significant associations of BMI with frailty measures. We investigated the associations of 29 markers with BMI in our cohort (Figure 5A). We observed seven inflammageing markers associated with BMI which included IL-6 and CRP. Therefore, we tested associations of frailty measures with inflammageing markers by controlling for confounding effects of BMI, age and sex. Based on BMI, we categorized participants into lean (less than 25), overweight (between 25.0 and 30) and obese (above 30). These analyses showed reduced numbers of associations for all frailty measures except the Rockwood Frailty Index (Figure 5B). The Frailty Index was associated with four, EQ-5D-3L with four instead of six, PF.SF36 with four instead of five and HG.SF36 with three instead of six inflammageing markers. IL-6 was not observed to be associated with PF.SF36 and CRP was only associated with HG.SF36 after correcting for BMI and age. This analysis clearly indicates the importance of considering the effect of BMI when interpreting the association between frailty measures and cytokine measurements.

**Figure 5:**
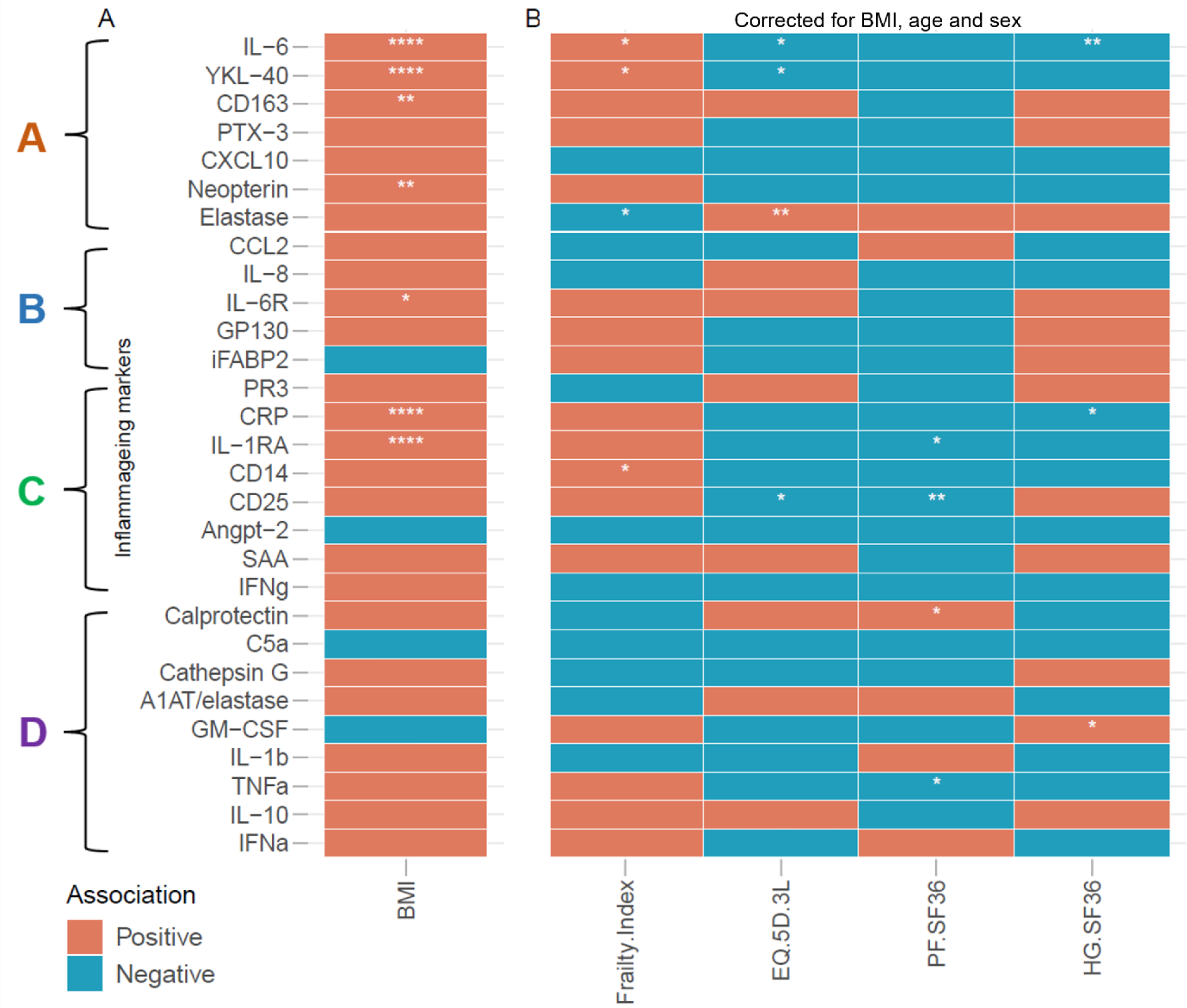
BMI has an important impact on levels of inflammageing markers and plays a role as a confounder in the association between frailty and inflammageing markers. In A, we show the associations of 29 markers with BMI in the VITAL cohort. In B, we show associations of 29 markers with frailty measures corrected for BMI, age and sex. The inflammageing markers are divided in groups A, B, C, and D based on their trajectory with age, see Figure 2. Statistical significance of the Spearman’s correlation coefficient is indicated with *: p<0.05, **: p<0.01, ***: p<0.001, ***: p<0.0001.

## DISCUSSION

In this comprehensive study we aimed to study the association between a large number of ageing-associated biomarkers with several measures of frailty. We showed that frailty is closely related to both age and BMI, requiring us to correct for these factors to truly understand the associations between the biomarkers and frailty. We found a number of markers, including IL-6, CRP, YKL-40, IL-1RA, PR3 and elastase to be associated with frailty measures. Our findings indicate that frailty is related to dysfunctional immunological processes that arise with ageing, such as low-grade inflammation and neutrophil dysfunction.

Ageing of the immune system is accompanied by increased levels of a wide range of cytokines and other pro-inflammatory markers in the blood. In this study we showed a positive association of age with 19 biomarkers. The underlying mechanisms may vary. Inflammageing is likely caused by increased recognition of danger signals and a reduced barrier function of the mucosa in the gut (2,3,35). Another cause of inflammageing is cellular senescence, which is a cellular fate caused by DNA damage resulting in cells that are prevented from proliferating, but characterized by a resistance to apoptosis and the production of a wide range of pro-inflammatory cytokines (36–38). Finally, BMI tends also to increase with age, and particularly a high fat mass associates with low-grade inflammation (33,34). The interplay between these and additional factors leads to a pattern of increasing inflammatory biomarkers with increasing age. Interestingly we observed that the increases with age seen for these biomarkers followed different patterns: some markers tend to increase continuously, some may increase at middle age but then plateau, and some increase only in the oldest adults. Further studies may reveal whether the distinct underlying causes of inflammageing may explain these different patterns.

The choice of frailty measures is important in studies that investigate the link between frailty and immunological processes. The four main frailty measures in this study likely reflect partially different types of frailty. The Frailty Index (29), focusses on deficits, which typically develop with age, and indeed the Frailty Index strongly correlated with age. The advantage of the Frailty Index is that it is a more objective instrument to measure frailty, rather than the EQ-5D-3L and SF-36 which were solely based on questionnaires. The associations of the Frailty Index with the biomarkers are less impacted by BMI compared those of the EQ-5D-3L and SF-36 domains, as all four associations between the Frailty Index and the biomarkers remained after correcting for BMI. The number of prescribed medications used correlated strongly with the other frailty measures, suggesting its potential use as an easily obtained and objective measure of frailty. Due to the use of home visits, The Frailty phenotype, by Fried *et al*. (2001), was not used in our analyses since it would require more space than generally available.

A number of markers that are typically associated with innate immunity activation were found to be associated with frailty measures. The pro-inflammatory cytokine IL-6 and the closely related acute-phase marker CRP were associated with frailty. IL-6 and CRP have often been proposed as potential markers of frailty (39–42). Also TNF-α levels have been found to associate with frailty, however this was not the case in this study, nor in another recent study (20). The association between IL-6/CRP levels and frailty may partly be explained by the close interplay with BMI. IL-6 and CRP levels have been associated with increased adiposity, and adipocytes are potent producers of IL-6, particularly during a state of chronic low-grade inflammation (32,43). In this study we show a strong relation between BMI and the frailty measures. When correcting for BMI, the association between IL-6 and three of the frailty measures remained.

We also included a number of markers that have less often been studied in the context of frailty. We found that YKL-40 levels associated with frailty measures, and even after correcting for age and BMI the association with the Frailty Index and EQ-5D-3L remained. YKL-40, (chitinase-3-like protein 1) is produced by certain types of neutrophils and macrophages and was previously found to play a role in orchestrating angiogenesis, tissue fibrosis, inflammation, oxidative tissue injury, and extracellular remodeling responses (44–49). As a biomarker that likely reflects immune activation and potentially peripheral macrophage or even microglial activation, YKL-40 has been associated with neurodegenerative diseases, including a role as a prognostic marker (50,51). Elevated YKL-40 levels have been also reported to be associated with atherosclerosis and cerebrovascular disease(52,53). In a study with 80-year old participants, YKL-40 levels associated with all-cause mortality, which also implies a link with frailty (54). Our results are in line with those previous observations.

The level of IL-1RA, an antagonist of pro-inflammatory IL-1 signaling, was also associated with age-corrected frailty, even though this relation was largely lost when we corrected for BMI. Indeed, IL-1RA levels have been associated with the risk for cardiovascular disease, which likely plays a prominent role in frailty (55). IL-1RA release is stimulated by inflammatory mediators including IL-1β and IL-6, in which is thought of as a negative feedback mechanism (56). Moreover, IL-1RA levels appeared to be closely associated with innate cells counts in the blood. Similarly, levels of neopterin, a marker of global immune activation described to increase with age, were associated with sex-corrected frailty, as previously described (23). However, this association did not persist when we corrected for age and/or BMI.

We also showed that two neutrophil-associated markers, PR3 and Elastase, associated negatively with frailty, despite their positive association with age. Both PR3 and Elastase are important proteases that neutrophils use to induce their extravasation from the blood and migration within the tissue (57). Elastase can be found in free form or bound to its inhibitor A1AT, but here only the free form of Elastase was associated with frailty. One study has investigated neutrophil dysfunction in the context of frailty (58). Here they discovered that elastase activity, measured indirectly via AαVal360 levels, was reduced in frail older adults. Interestingly, elastase levels in this study and AαVal360 levels in the study by Wilson *et al*. do increase with age, but also associate negatively with frailty. Potentially, the dysfunctional neutrophils contribute to the delayed disposal of pathogens, resulting in lower thresholds for inflammatory responses.

An additional finding in this study was the observed association of serum angiopoietin-2 levels with both anti-CMV and EBV seropositivity. The association of angiopoietin-2 levels with chronic CMV and EBV infections has not been reported before, but increased angiopoietin-2 levels have been related to chronic hepatitis C infection (59). Angiopoietin-2 is an inflammation-associated protein that promotes angiogenesis, the formation of new small blood vessels (60). Angiogenesis is instigated by angiopoietin-2 by disrupting the homeostatic angiopoietin-1 – Tie2 signaling in the presence of vascular endothelial growth factor (VEGF). Potentially, the variety of liver manifestations that are associated with CMV and EBV infection explain the higher angiopoietin levels; a comparable mechanism was observed in patients with chronic hepatitis C (59,61). In addition to angiopoietin-2, we also found an association of macrophage activation marker sCD163 with anti-CMV seropositivity, confirming a prior report (62), and monocyte chemoattractant CCL2 with anti-EBV antibodies. We considered the CMV and EBV status of the participants due to the link with immune ageing (6,30), however we found no additional overlap in markers associated with frailty and markers associated with CMV and EBV status.

Strengths of this study include the use of a broad prospective cohort with participants from a large age range (25-92 years old). Additionally, we used both objective and subjective determinants of frailty and focus on a comprehensive set of markers that were associated with mechanisms behind inflammageing. Frailty associations with inflammageing markers were performed with sophisticated analyses in which we corrected for the confounders age, sex and BMI. Weaknesses include the fact that male participants in this study were on average older than the female participants. This confounder, was however corrected for in most on the analyses.

## CONCLUSION

We here show that several inflammageing markers are independently associated with frailty, even when corrected for age, sex and BMI. These associations provide clues on reasons why ageing affects people differently with regards to their physical health and resilience against stressors. These biomarkers could potentially be employed in ageing studies to assess frailty instead or in addition to the Frailty Index or questionnaires, that are both time consuming and less consistent between different studies. The immunological mechanisms and cellular source of these frailty-associated biomarkers require further studies to validate the observed markers herein in a global inflammageing signature. Thus, it would be important to associate these biomarkers to extensive immunophenotyping of the same participants at the same visit. The main goal of the VITAL cohort concerns the investigation of the vaccine responses in the ageing population. Therefore, future studies will aim to associate both frailty measures and inflammageing markers with outcomes of vaccination.

## MATERIALS AND METHODS

### The VITAL cohort

The VITAL cohort consists of individuals aged 25-90. Younger (age 25-49) and middle-aged (age 50-64) participants were employees recruited from Health institutes at the Utrecht Science Park and Spaarne Hospital (Hoofddorp). The older participants (age >65) were recruited from a previous study on influenza in older adults (25,26). We excluded individuals with previous adverse reactions to vaccination or for whom blood drawing may be harmful. We also excluded immunocompromised individuals due to disease, or immune-mediating medication, such as high dose glucocorticoids or chemotherapy during the study or in the previous three years. Also, participants were excluded if they did not receive an influenza vaccination in the 2018-2019 season. Participants were recruited for the VITAL longitudinal intervention cohort which aims aim at getting a better insight of the influence of age and age-related changes on vaccine-induced immune responses, and at gaining knowledge on the trajectory of immune decline in older adults and middle-aged adults in comparison to adults (18,19).

Here, we included clinical data, questionnaires and serum and plasma samples from 317 participants at the first study visit in 2019, which was prior to influenza vaccination. Blood was drawn from each participant; serum and EDTA-plasma were stored within 8 hours at -80°C until further use.

### Assessment of frailty status in the VITAL cohort

As our main determinant for frailty, we designed a Frailty Index for the VITAL cohort, following the criteria that were described by Rockwood and Minitski (27). The deficits that are included in the Frailty Index should meet defined criteria. These criteria state that: each deficit counts equally in the Frailty Index; the included deficits must cover a range of functions; deficits are associated with health; the prevalence of the deficit increases with age; the parameter (whether or not someone has the deficit) is measured in ≥95% of the study population; the deficit is present in ≥1% of the study population; and the deficit does not occur in >80% of the study population <80 years (13,14,27–29). The Frailty Index in the VITAL cohort contains 31 deficits, which is within the recommended range of 30-40 deficits (see Supplementary Table S1 for the full list of deficits and scoring methods). Deficits included in the Frailty Index covered a range of systems, including cardiovascular, gastrointestinal, sensory, motor and neural systems. If participants were missing data for ≥20% of the deficits, the Frailty Index could not be determined. The Frailty Index results in a score ranging from 0 (least frail) to 1 (most frail), although scores above 0.67 have been reported to be too high to be compatible with life.

Two additional methods for defining frailty were based on self-reported questionnaires, comprising the EQ-5D-3L and the SF36. The EQ-5D-3L is a questionnaire on five dimensions: mobility, self-care, usual activities, pain/discomfort and anxiety/depression (16). For each dimension, the participants scored their health on three levels. Using the population norms of the Netherlands, a composite score ranging from -1 (most frail) to 1 (least frail) was set. The SF-36, a set of generic, coherent, and easily administered quality-of-life measures, comprises 36 multiple-choice questions (15). The complete SF-36 questionnaire results in scores in eight domains, ranging from 0 (most frail) to 100 (least frail). In this study, we only used data from two domains, PF and HG, as they displayed the most variation in scores. Additionally, we compared the four measures of frailty with the Health Rating score and the number of prescription medications. The Health Rating score, ranging from 0 (most frail) to 10 (least frail) was a simple rating scored by the participant based on the question “I rate my health with a score of …”. Prescription medications that are not indicative of a frail state (contraception, malaria prophylaxis, and one-time sedation for medical procedures) were not included in the number of medications.

### Laboratory measurements

Levels of biomarkers reflecting inflammageing were measured with different techniques in serum or plasma samples of the VITAL participants (Supplementary Table S2). Twenty biomarkers were measured at the University Medical Center Groningen (UMCG) in the Netherlands, using either Luminex multiplex assays or ELISA assays. An additional nine markers were assessed in Cimi-Paris (France), by ELISA or by the high-sensitivity Quanterix assays (Simoa or Corplex). For technical information, including dilutions and standard curve ranges, see Supplementary Table S2. Samples that were above or below the range of the standard curve were set at a fixed value, as indicated in Supplementary Table S2. Assays were performed in accordance with the manufacturer’s recommendations. The absolute number of monocytes and neutrophils was assessed by the DxH 500 Hematology Analyzer (Beckman Coulter).

Tests to determine CMV and EBV status at the first visit, based on immunoglobulin G antibodies against CMV and EBV, was performed by multiplex immunoassay (MIA), as described before (30,31). For CMV, a concentration of <4 relative units (RU) ml−1 was categorized as seronegative, ≥4 and <7.5 RU ml−1 as borderline, and ≥7.5 RU ml−1 as seropositive. For EBV, a concentration of <16 RU ml−1 was categorized as seronegative, ≥16 AU ml−1, and <22 RU ml−1 as borderline and ≥ 22 RU ml−1 as seropositive.

### Statistics

Statistical analysis and data visualization were done in R (v4.2.1) statistical programming language. Associations between two numeric variables (frailty vs age and frailty vs cytokine measurements) were tested with the permutational framework for Spearman’s test implemented in the coin (v1.4.2) R package. The analysis considered sex as a ‘block’ for stratification and the null distribution of the test statistic was approximated via Monte-Carlo resampling (nresample=100000). The associations of the different frailty measures with cytokines in older adults included two different approaches. First comparison tested associations without correcting for sex and age and second comparison included stratification combined for age and sex to avoid cofounding in this study.

Additionally, multiple regression analyses were performed in IBM SPSS Statistics 28 with the Frailty Index scores as dependable outcome and age and the biomarkers as predictable variables. Due to the non-normal distribution of the data, the Frailty Index scores and biomarker levels were log transformed. Participants with missing data were also excluded from the analysis. This analysis was performed for all participants, and separately for female and male participants. Multicollinearity was checked with VIF statistics < 10 and thereafter the collinearity diagnostics.

The PCA was performed using the ggplot2 package in RStudio 2022.07.2. The same log-transformed biomarkers were used as in the regression analysis, and only participants above 60 years of age with complete data for all markers were included. The amount of variance explained by Principle Components (PC) 1 and 2 are mentioned on the axis.

## DECLARATIONS

### Ethics approval and consent to participate

Approval from the institutional review board for this study was obtained from the Medical Research Ethics Committee Utrecht (EudraCT Number: 2019–000836–24) and all participants signed for informed consent. All procedures were in accordance with the declaration of Helsinki.

### Consent for publication

Not applicable

### Availability of data and materials

The datasets analysed during the current study are available from the corresponding author on reasonable request.

### Authors’ contributions

JvB, AMB and DvB conceptualized the clinical study. YvS, MCAV, NGM collected the data. YvS, SAS and MvdH performed the statistical analysis. DvB and DS supervised the study. YVS wrote the original draft. SAS, MvdH, AMB, JvB, DvB, EJMT and DS were involved in writing, review and editing. All authors read and approved the final manuscript.

## Acknowledgements

The VITAL project has received funding from the Innovative Medicines Initiative 2 Joint Undertaking (JU) under grant agreement No. 806776 and the Dutch Ministry of Health, Welfare and Sport. The JU receives support from the European Union’s Horizon 2020 research and innovation programme. The authors thank Wivine Burny, Carole Desion, Roberto Zoffoli and Daniel Laroque, partners in the VITAL consortium, for their feedback on this study. We also thank the volunteers for participating in this study.

## Additional file 1

Frailty manuscript supplementary data V5.pdf

Supplementary data: contains supplementary tables 1-2 and supplementary figures 1-3.

