## Supplementary data for "Frailty is related to serum inflammageing markers: results from the VITAL study"

**Supplementary Table S1: Full list of the deficits and their scoring as included in the frailty index developed in this study.**

| Deficit | Questions/measurements included in deficit | Scoring | |
|---|---|---|---|
| High blood pressure | - “Have you ever been found to have high blood pressure?”<br>- Systolic blood pressure measurement (mmHg) (measured twice)<br>- Diastolic blood pressure measurement (mmHg) (measured twice) | - “yes”<br>- Participant ≤70 year + both systolic measurements ≥140 mmHg<br>- Participant >70 year + both systolic measurements ≥150 mmHg<br>- Both diastolic measurements ≥80 mmHg<br>- Participant ≤70 year + at least one systolic measurements <140 mmHg + at least one diastolic measurement <80 mmHg<br>- Participant >70 year + at least one systolic measurements <150 mmHg + at least one diastolic measurement <80 mmHg | 1<br>1<br>1<br>1<br>0<br>0 |
| Cardiovascular disease | - “Have you ever had a heart attack/myocardial infarction?”<br>- “Have you ever undergone heart bypass surgery?”<br>- “Have you ever undergone coronary angioplasty/percutaneous coronary intervention?”<br>- “Have you ever undergone heart catheterization?”<br>- “Have you ever undergone a vascular surgery?”<br>- “Have you ever ended up in the hospital because of heart failure?”<br>- If a participant indicated, at any other question, that they had any form of heart or vessel deficit that was not covered by one of the above questions | - “Yes” to any<br>- “No” to all | 1<br>0 |
| Stroke | - “Have you ever had a stroke?” | - “Yes”<br>- “No” | 1<br>0 |
| Arrhythmia | - “Have you ever undergone a pacemaker or ICD surgery?”<br>- If a participant indicated, at any other question, that they had any form of arrhythmia that was not covered by the above question | - “Yes” to any<br>- “No” to both | 1<br>0 |
| Hypercholesterolemia | - LDL values measured from participants peripheral blood (mmol/L)<br>- If a participant indicated, at any question, that they had hypercholesterolemia | - Participant indicated that they had hypercholesterolemia<br>- LDL ≥2.6 mmol/L<br>- None of the above | 1<br>1<br>0 |
| Grip strength (GS) | - Grip strength measurement (kg) (measured three times) | - Participant is a man with:<br> - BMI≤24 + average GS ≤29 kg<br> - BMI 24.1-28 + average GS ≤30 kg<br> - BMI>28 + average GS ≤32 kg<br>- Participant is a woman with:<br> - BMI≤23 + average GS ≤17 kg<br> - BMI 23.1-26 + average GS ≤17.3 kg<br> - BMI 26.1-29 + average GS ≤18 kg<br> - BMI>29 + average GS ≤21 kg<br>- None of the above | 1<br>1<br>0 |
| Physiotherapist | - “Do you receive care/treatment from a physiotherapist?” | - “Yes”<br>- “No” | 1<br>0 |
| Psychiatrist | - “Are you under treatment with a psychiatrist?” | - “Yes”<br>- “No” | 1<br>0 |

| | | | |
|---|---|---|---|
| BMI | - BMI (kg/m$^2$) calculated from participants height and weight | - BMI <18.5 or ≥30 kg/m$^2$<br>- BMI ≥25 and >30 kg/m$^2$<br>- None of the above | 1<br>0.5<br>0 |
| Appetite | - "My appetite is:" | - "Bad"<br>- "Mediocre"<br>- "Average"<br>- "Good" | 1<br>0.5<br>0<br>0 |
| Dietician | - "Do you receive care/treatment from a dietician?" | - "Yes"<br>- "No" | 1<br>0 |
| Caregiver | - "Do you receive informal care from a caregiver?"<br>- "Do you receive care/treatment from a home care worker?"<br>- "Do you receive care/treatment from a district nurse?" | - "Yes" to any<br>- "No" to all | 1<br>0 |
| Treatment from a general health care worker | - "Are you under treatment with a general practitioner (GP)?"<br>- "Do you receive care/treatment from a nurse in your general practice?" | - "Yes" to any<br>- "No" to both | 1<br>0 |
| Hearing | - "Can you have a conversation in a group of 3 or more people (with hearing aid if necessary)?" | - "No, I cannot"<br>- "Yes, with a lot of effort"<br>- "Yes, with a little effort"<br>- "Yes, without effort" | 1<br>0.66<br>0.33<br>0 |
| Seeing | - "Are your eyes good enough to be able to read the small letters in the newspaper (with glasses or contact lenses if necessary)?" | - "No, I cannot"<br>- "Yes, with a lot of effort"<br>- "Yes, with a little effort"<br>- "Yes, without effort" | 1<br>0.66<br>0.33<br>0 |
| Cancer | - "Do/did you have a type of cancer?" | - "Yes"<br>- "No" | 1<br>0 |
| Diabetes | - "Have you been diagnosed with diabetes?" | - "Yes"<br>- "No" | 1<br>0 |
| Bowel disorder | - "Do you have severe or persistent bowel disorders longer than 3 months?" | - "Yes, diagnosed by a doctor"<br>- "Yes, not diagnosed by a doctor"<br>- "No" | 1<br>1<br>0 |
| Thyroid issues | - "Do you have thyroid issues?" | - "Yes, diagnosed by a doctor"<br>- "No" | 1<br>0 |
| Internist | - "Are you under treatment with an internist?" | - "Yes"<br>- "No" | 1<br>0 |
| Disease of the nervous system | - "Do you have a disease of the nervous system (Parkinson's, MS, epilepsy)?" | - "Yes, diagnosed by a doctor"<br>- "Yes, not diagnosed by a doctor"<br>- "No" | 1<br>1<br>0 |
| Dizziness | - "Do you have dizziness with falling?" | - "Yes, diagnosed by a doctor"<br>- "Yes, not diagnosed by a doctor"<br>- "No" | 1<br>1<br>0 |
| Joint inflammation | - "Do you have chronic joint inflammation (inflammatory rheumatism, chronic rheumatism, rheumatoid arthritis)?" | - "Yes, diagnosed by a doctor"<br>- "Yes, not diagnosed by a doctor"<br>- "No" | 1<br>1<br>0 |
| Bone calcification | - "Do you have bone calcification (osteoporosis)?" | - "Yes, diagnosed by a doctor"<br>- "Yes, not diagnosed by a doctor"<br>- "No" | 1<br>1<br>0 |
| Back problems | - "Do you have severe or persistent back problems (including hernia)?" | - "Yes, diagnosed by a doctor"<br>- "Yes, not diagnosed by a doctor"<br>- "No" | 1<br>1<br>0 |
| Urine loss | - "Do you have involuntary urine loss (incontinence)?" | - "Yes, diagnosed by a doctor"<br>- "Yes, not diagnosed by a doctor"<br>- "No" | 1<br>1<br>0 |
| Shingles | - "Have you ever experienced shingles?" | - "Yes"<br>- "No" | 1<br>0 |
| Lung disease | - "Do you have a lung disease?" | - "Yes, diagnosed by a doctor"<br>- "Yes, not diagnosed by a doctor"<br>- "No" | 1<br>1<br>0 |
| Respiratory tract infection | - "In the past 12 months, how many times have you had an upper respiratory tract infection for which you have visited the | - Total number of visits in the past 12 months is ≥2<br>- Total number of visits in the past 12 months is 1 | 1<br>0.5 |

| | | | |
|---|---|---|---|
| | doctor? (cold with fever, ear infection, throat infection, nasal sinus infection)"<br>- "In the past 12 months, how often have you had a lower respiratory tract infection for which you have visited the doctor? (bronchitis, pneumonia)" | - Participant did not visit the doctor for a upper or lower respiratory tract infection in the past 12 months | 0 |
| Surgeon/orthopedist | - "Are you under treatment with a surgeon/orthopedist?" | - "Yes"<br>- "No" | 1<br>0 |
| Medication usage | - "Have you used prescription medication in the 3 months prior to the study? and/or Have you used prescription medication during the study?" | - Participant used ≥5 medications<br>- Participant used <5 medications | 1<br>0 |

Deficits were scored according to the example and guidelines of Rockwood and Mitnitski, and cut points/normal values for blood pressure and LDL values were adapted from the Dutch General Practitioners Association. Medications that were not indicative of a frail state (contraception, malaria prophylaxis, and one-time sedation for medical procedures) were not included in the "Medication usage" deficit. Questions and answers between quotation marks were the literal, translated, questions asked to participants, and answers given by participants.

**Supplementary table S2: technical information on each assay used to obtain biomarker concentrations**

| Marker | Method | Lab | Material | Dilution | Lowest limit | Highest limit | Unit | Company | Missing values | Samples below detection set at | Samples above detection set at |
|---|---|---|---|---|---|---|---|---|---|---|---|
| Angiopoietin-2 | Luminex | UMCG | serum | 4 | 270 | 65720 | pg/mL | R&D | 4 | | |
| C5a | Luminex | UMCG | serum | 4 | 3.03 | 735.56 | ng/mL | R&D | 4 | 1 | |
| CCL2 | Luminex | UMCG | serum | 4 | 123 | 29880 | pg/mL | R&D | 4 | | |
| sCD25 | Luminex | UMCG | serum | 4 | 107 | 26000 | pg/mL | R&D | 4 | | |
| sCD163 | Luminex | UMCG | serum | 4 | 22.0 | 5346 | pg/mL | R&D | 4 | | |
| CXCL10 | Luminex | UMCG | serum | 4 | 4.12 | 1000 | pg/mL | R&D | 4 | | |
| sGP130 | Luminex | UMCG | serum | 4 | 1.85 | 450 | ng/mL | R&D | 4 | | |
| IL-1RA | Luminex | UMCG | serum | 4 | 117 | 28480 | pg/mL | R&D | 4 | | |
| sIL-6R | Luminex | UMCG | serum | 4 | 0.43 | 103 | ng/mL | R&D | 4 | | |
| PTX-3 | Luminex | UMCG | serum | 4 | 658 | 159880 | pg/mL | R&D | 4 | | |
| YKL-40 | Luminex | UMCG | serum | 4 | 1.76 | 429 | ng/mL | R&D | 4 | | |
| CRP | Luminex | UMCG | serum | 400 | 40.8 | 9920 | ng/mL | R&D | 7 | | 15000 |
| sCD14 | Luminex | UMCG | serum | 400 | 90.6 | 22012 | ng/mL | R&D | 7 | | |
| IL-8 | ELISA | UMCG | serum | 1 | 1 | 64 | pg/mL | R&D | 3 | | 100 |
| Calprotectin | ELISA | UMCG | serum | 200 | 330 | 20000 | ng/mL | Hycult | 3 | | |
| SAA | ELISA | UMCG | serum | 200 | 0.01 | 43.8 | ug/mL | in-house* | 3 | 0.005 | |
| Elastase | ELISA | UMCG | plasma | 40 | 16 | 1000 | ng/mL | Hycult | 11 | | 1500 |
| PR3 | ELISA | UMCG | plasma | 10 | 6.30 | 400 | ng/mL | Hycult | 11 | 5 | 600 |
| Cathepsin G | ELISA | UMCG | plasma | 20 | 1.25 | 40 | ng/mL | Hycult | 28 | 1 | |
| A1AT-Elastase | ELISA | UMCG | plasma | 4000 | 6.24 | 4000 | ng/mL | Hycult | 37 | | |
| Neopterin | ELISA | INSERM | serum | 1 | 1.35 | 111 | nmol/l | Tecan | 11 | | |
| iFABP2 | ELISA | INSERM | serum | 5 | 0 | 1000 | pg/ml | R&D | 11 | | |
| GM-CSF | Simoa HD-1 | INSERM | serum | 4 | 0 | 120 | pg/ml | Quanterix | 11 | 0 | |
| IFN-$\alpha$ | Simoa HD-1 | INSERM | serum | 2 | 0 | 60 | pg/ml | Quanterix | 11 | 0 | |
| IL-1$\beta$ | Corplex | INSERM | serum | 4 | 0 | 100 | pg/ml | Quanterix | 11 | 0 | |
| TNF-$\alpha$ | Corplex | INSERM | serum | 4 | 0 | 400 | pg/ml | Quanterix | 11 | | |
| IFN-γ | Corplex | INSERM | serum | 4 | 0.0004 | 100 | pg/ml | Quanterix | 11 | | |
| IL-6 | Corplex | INSERM | serum | 4 | 0.0001 | 600 | pg/ml | Quanterix | 11 | | |
| IL-10 | Corplex | INSERM | serum | 4 | 0 | 100 | pg/ml | Quanterix | 11 | 0 | |

ELISA: Enzyme Linked Immune Sorbent Assay, UMCG: University Medical Center Groningen (Groningen, the Netherlands), Cimi-Paris: Centre d'immunologie et des maladies infectieuses (Paris, France)*: SAA was measured according to the protocol previously published (Hazenberg BP et al, 1999, Ann Rheum Dis 1999;58(2):96-102). Missing values were due to unavailable plasma samples (e.g. Elastase), insufficient serum volume (e.g. Cathepsin G), or due to technical errors (e.g. CCL2).

**A**

| | Frailty Index | EQ- 5D-3L | SF-36 PF | SF-36 HG | Health Rating |
|---|---|---|---|---|---|
| EQ-5D-3L | -0.47** | | | | |
| SF-36 PF | -0.57** | 0.56** | | | |
| SF-36 HG | -0.47** | 0.47** | 0.50** | | |
| Health Rating | -0.42** | 0.50** | 0.46** | 0.64** | |
| # of medications | 0.70** | -0.31** | -0.45** | -0.41** | -0.30** |

**B**

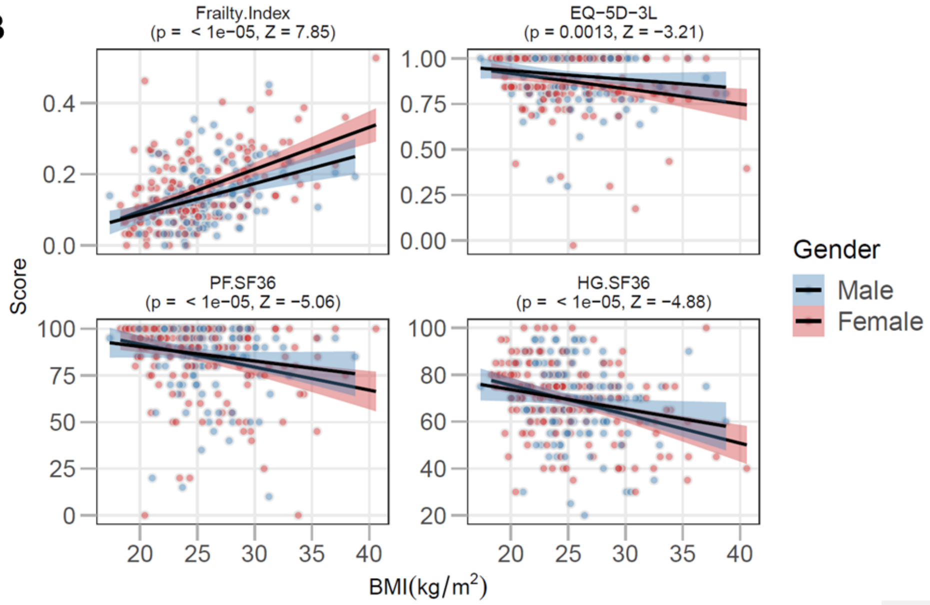


**Supplementary Figure S1: correlations of frailty measures with each other and BMI.** In A, Spearman correlation coefficients and cell colors are shown indicating the strength and direction of the associations. **: p<0.01. In B, correlations of each frailty measure with BMI are shown. Strength and significance of the sex-corrected Spearman correlation coefficient is indicated in each graph.

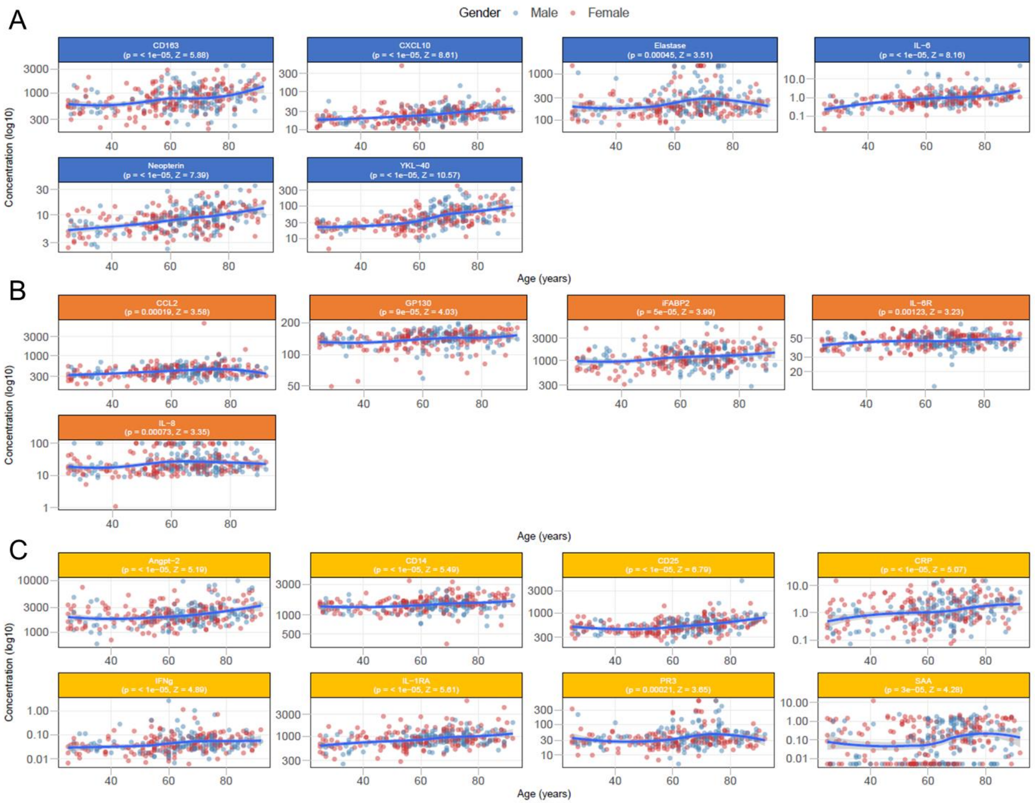


**Supplementary Figure S2: Associations of all 29 biomarkers with age.** Shown are individual values and moving medians (IQR) for each biomarker. Biomarkers marked with a * showed a significant association with age (p<0.05).

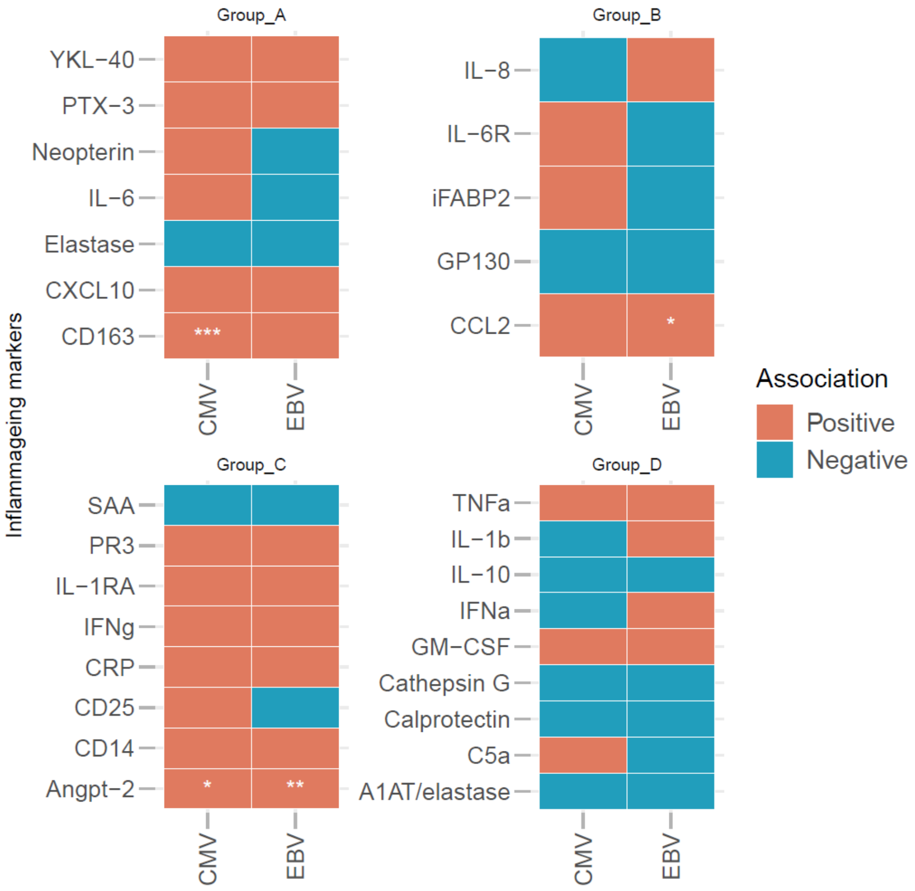


**Supplementary Figure S3: associations of inflammageing markers with CMV and EBV status.** Shown are sex-corrected associations of the biomarkers with CMV- and EBV-status. Angiopoietin-2 levels were significantly higher in participants that were seropositive for CMV- and EBV antibodies compared to participants that were seronegative. A similar association of sCD163 levels was found with CMV status, and CCL2 levels with EBV status. Statistical testing was done with the Wilcoxon signed rank test, '+' indicates statistical significance ($p<0.05$). CMV: cytomegalovirus, EBV: Epstein-Barr virus.

**Met opmerkingen [d1]:** In the Figure we need to add s for CD163→ sCD163

**Met opmerkingen [d2]:** *p<0.05
** p< 0.01???
